# Regulation of intercellular TARGET OF MONOPTEROS 7 protein transport in the *Arabidopsis* root

**DOI:** 10.1101/122960

**Authors:** Kuan-Ju Lu, Bert De Rybel, Hilda van Mourik, Dolf Weijers

**Affiliations:** Laboratory of Biochemistry, Wageningen University, Stippeneng 4, 6708 WE, Wageningen, The Netherlands.; Ghent University, Department of Plant Biotechnology and Bioinformatics, Technologiepark 927, 9052 Ghent, Belgium; VIB Center for Plant Systems Biology, Technologiepark 927, 9052 Ghent, Belgium

**Author notes:** Present address: Laboratory of Molecular Biology, Wageningen University, 6708PB, Wageningen, The Netherlands (Hilda van Mourik). Corresponding author: Dolf Weijers.

**Keywords:** TMO7, Plasmodesmata, cell-cell communication, protein transport, embryogenesis, RAM

## Abstract

Intercellular communication coordinates hypophysis establishment in the Arabidopsis embryo. Previously, TARGET OF MONOPTEROS 7 (TMO7) was reported to be transported to the hypophysis, the founder cell of the root cap, and RNA suppression experiment implicated its function in embryonic root development. However, it remained unclear what protein properties and mechanisms mediate TMO7 protein transport, and what role the movement plays in development. Here, we report that in the post-embryonic root, TMO7 and its close relatives are transported into the root cap through plasmodesmata in a sequence, but not size dependent manner. We also show that nuclear residence is critical for TMO7 transport, and postulate that modification, potentially phosphorylation, labels TMO7 for transport. Additionally, three novel CRISPR/Cas9-induced *tmo7* alleles confirmed a role in hypophysis division, but suggest complex redundancies with close relatives in root formation. Finally, we demonstrate that TMO7 transport is biologically meaningful, as local expression partially restores hypophysis division in a plasmodesmatal protein transport mutant. Our study identifies motifs and amino acids critical for TMO7 protein transport and establishes the importance of TMO7 in hypophysis and root development.

**Summary Statement:** Unique protein motifs, subcellular localization and post-translational modification, rather than protein size regulate plasmodesmatal transport of TMO7 family proteins during Arabidopsis root development.

## Introduction

Intercellular communication is vital for development of multi-cellular organisms (Long et al., 2015), for example to coordinate distant events or locally organize tissues (Balkunde et al., 2017; Corbesier et al., 2007; Kurata et al., 2005; Nakajima et al., 2001; Pi et al., 2015; Sessions et al., 2000; Tamaki et al., 2007; Yadav et al., 2011). Plant cell walls provide robust physical support to build elaborate tissues and structures; however, cell walls also hamper cell migration, and it is therefore important for plant cells to receive positional cues from their surroundings for proper development (ten Hove et al., 2015). To overcome the physical barrier that cell walls pose to intercellular communication, plants evolved specialized channels connecting neighbouring cells, named plasmodesmata (PD). PD contain compressed ER tubes, allowing transport of molecules between cells (Otero et al., 2016).

Many plant tissue patterning events have been shown to involve cell-cell communication (Bernhardt et al., 2005; Helariutta et al., 2000; Kim et al., 2003; Kurata et al., 2005; Rodriguez et al., 2016; Schlereth et al., 2010). One of the earliest of such events in plant ontogeny occurs during early embryogenesis in the flowering plant *Arabidopsis thaliana* (ten Hove et al., 2015). Formation of the embryonic root requires the AUXIN RESPONSE FACTOR 5/MONOPTEROS (ARF5/MP) transcription factor, and *mp* mutants fail to form a root (Berleth and Jurgens, 1993; Hardtke and Berleth, 1998). The root forms from two cell populations: a set of embryonic cells and their extra-embryonic neighbour (the hypophysis) (Scheres et al., 1994). Both these cell populations develop abnormally in the *mp* mutant (Berleth and Jurgens, 1993; Hardtke and Berleth, 1998), but expression of MP only in the embryonic cells complements not only the embryonic but also the hypophysis defect, indicating a non-cell-autonomous function(Weijers et al., 2006). Immobility of MP protein suggests the existence of downstream mobile signals (Weijers et al., 2006). A direct MP downstream target, TARGET OF MONOPTEROS 7 (TMO7), is transcribed in embryonic cells but the protein is found in the neighbouring hypophysis, strongly suggesting protein transport. Fusing TMO7 to triple GFP (3xGFP) prevented protein accumulation in the hypophysis, suggesting that a size restriction to movement (Schlereth et al., 2010). Based on RNA suppression approaches and local expression, TMO7 appears to contribute either to establishing hypophysis identity or to controlling its cell division plane (Rademacher et al., 2012; Schlereth et al., 2010). TMO7 is an atypical basic Helix-Loop-Helix (bHLH) transcription factor that lacks the basic region, and is significantly smaller than most mobile transcription factors studied to date. Key questions are how this protein is transported, what regulates transport, and what role this plays in development.

Several transcription factors that control plant development have been shown to move between cells. The *Zea mays KNOTTED* 1 gene (*KN1*), and its homolog in *Arabidopsis thaliana, SHOOT MERISTEMLESS* (*STM*) encode homeobox-domain (HD) proteins which help maintain the shoot apical meristem (SAM) cells undifferentiated, and move directionally from inner to outer cell layers (Kim et al., 2003; Long et al., 1996; Lucas et al., 1995; Vollbrecht et al., 1991). The LEAFY (LFY) protein, a helix-turn-helix transcription factor that also participates in SAM development, appears to move by non-directional diffusion (Sessions et al., 2000; Wu et al., 2003). In addition, WUSCHEL (WUS), another key HD-containing transcription factor that regulates and maintains SAM activity, was also reported to move through PD (Daum et al., 2014; Yadav et al., 2011). Likewise, in root development there are several known mobile transcription factors, among which the SHORTROOT protein has been studied in most detail. *SHORTROOT* (*SHR*), a GRAS family transcription factor, is transcribed in stele tissues, but the SHR protein subsequently moves a layer outward to the endodermis and quiescent centre (QC) cells *SCARECROW* (*SCR*), another protein in GRAS family, is expressed (Helariutta et al., 2000; Nakajima et al., 2001). Together with SCR, SHR regulates the expression of C2H2 zinc-finger domain transcription factors, *JACKDAW* (*JKD*) and *MAGPIE* (*MGP*) in the endodermis, which forms a feed forward loop with *SCR* (Welch et al., 2007). SHR localizes in both cytoplasm and nucleus at stele; the interaction with SCR, JKD and MGP after movement, however, relocates the protein complex exclusively to the nucleus and prohibits the further movement of SHR (Gallagher et al., 2004; Nakajima et al., 2001; Welch et al., 2007). This transport regulation is critical for proper pattern formation of roots, as ectopic expression of SHR by the ubiquitous 35S promoter alters cell fate and creates multiplication of cell layers in root (Nakajima et al., 2001). Despite the importance of intercellular protein transport, the underlying mechanism is still largely elusive.

To reveal protein transport mechanisms, efforts have focused on identifying the critical transport elements in mobile transcription factors. However, the protein domains that mediate transport of different proteins do not seem to have common features (Gallagher et al., 2014), and it is therefore unclear how mobile proteins are selected for transport. Currently, the accumulation and degradation of callose, a β-1,3-glucan, at the neck region of the PD aperture by Callose Synthase (CalS) and β-1,3-glucanase, respectively, is the most prominent mechanism regulating cell-cell communication (Burch-Smith and Zambryski, 2012; Gallagher et al., 2014). A dominant *CalS* mutant, *cals3-2d*, was identified to regulate the accumulation of callose in Arabidopsis. By constructing a inducible iCalSm system, transport of SHR and its regulatory microRNA (miR165/166) was demonstrated to be blocked by the accumulation of callose at PD (Vaten et al., 2011). Yet, the accumulation of callose regulates the dilation of PD by physically closure and would not be expected to contribute to selectivity. Thus, the molecular mechanism of selective protein transport is still not well understood.

Here, we report that the TMO7 protein moves through PD, and that its movement contributes to hypophysis division. We show that TMO7 protein mobility is shared with a small set of TMO7-like proteins. We further show that sequence, not protein size determines mobility, and identified protein motifs critical for subcellular localization and transport. Our study provides a framework for understanding selective transport of transcription factors in the Arabidopsis root.

## Results

### TMO7 moves through plasmodesmata in the Arabidopsis root

TMO7 is expressed in the early Arabidopsis embryo, is transported from the pro-embryo to the hypophysis, and RNA suppression interferes with embryonic root formation (Schlereth et al., 2010). However, expression levels are extremely low and immunofluorescence is required to visualise the TMO7-GFP fusion protein in the embryo. *TMO7* was originally identified as a MP/BDL-dependent gene in a transcriptome study on seedlings (Schlereth et al., 2010), and we therefore addressed whether the post-embryonic root would represent a more accessible model for studying protein movement. We first observed the expression pattern of p*TMO7*::n3GFP (nucleus localization signal - 3 times GREEN FLUORESCENT PROTEIN), and compared it to p*TMO7*::TMO7-GFP, and p*TMO7*::TMO7-3GFP in root tips (Fig. 1, Schlereth et al., 2010). The *TMO7* promoter is expressed in meristematic and lateral root cap cells surrounding the QC and columella cells with a shootward-declining gradient. While nearly absent, very weak expression can also be observed in columella cells (Fig. 1A, B). In p*TMO7*::TMO7-GFP plants, we observed strong fluorescent signals not only in cells with high promoter activity, but also in QC and columella cells (referred as columella cells hereafter), indicating the rootward movement of TMO7-GFP protein, consistent with the reported movement of TMO7 in the Arabidopsis embryo (Fig. 1C; Schlereth et al., 2010). In contrast to TMO7-GFP, the expression of TMO7-3GFP is highly correlated with the *TMO7* promoter activity (Fig. 1B, D). Interestingly, the TMO7-3GFP protein seems to form aggregates in several cells, especially in cells above the QC (Fig. 1D). Also, the TMO7-3GFP protein is mostly excluded from the nucleus as low or no fluorescence signal was detected in the nucleus (Fig. 1D). The absence of fluorescence signal in columella cells indicates that movement of TMO7-3GFP is disrupted; however, weak GFP fluorescence was detected in cells below the QC in p*TMO7::*TMO7-3GFP roots (Fig. 1D, asterisk mark), which might be due to the weak expression of the *TMO7* promoter. To distinguish the weak activity of the *TMO7* promoter in columella cells from the signal derived from protein transport, the per-pixel fluorescence intensity ratio between columella cells and the rest of meristematic region was quantified (Fig. 1F and Fig. S1, see detailed description in Experimental Procedures). Consistent with the qualitative observation, the fluorescence intensity in columella cells is 28.8 ± 5.5% (mean ± SD, n=19) of the fluorescence in the meristematic region in p*TMO7::*n3GFP lines. The ratios in p*TMO7::*TMO7-GFP and p*TMO7::*TMO7-3GFP are 76.9 ± 9.6% and 47.5 ± 7.5% (n=19 and 18; mean±SD), respectively (Fig. 1F). These results indicate that the increase of fluorescence in columella cells in p*TMO7::*TMO7-GFP is due to the transport of TMO7-GFP, which is largely prevented in p*TMO7::*TMO7-3GFP. Importantly, these results show that, as in embryos, TMO7 protein is mobile in the post-embryonic root, and that it involves similar constraints.

**Fig 1.**
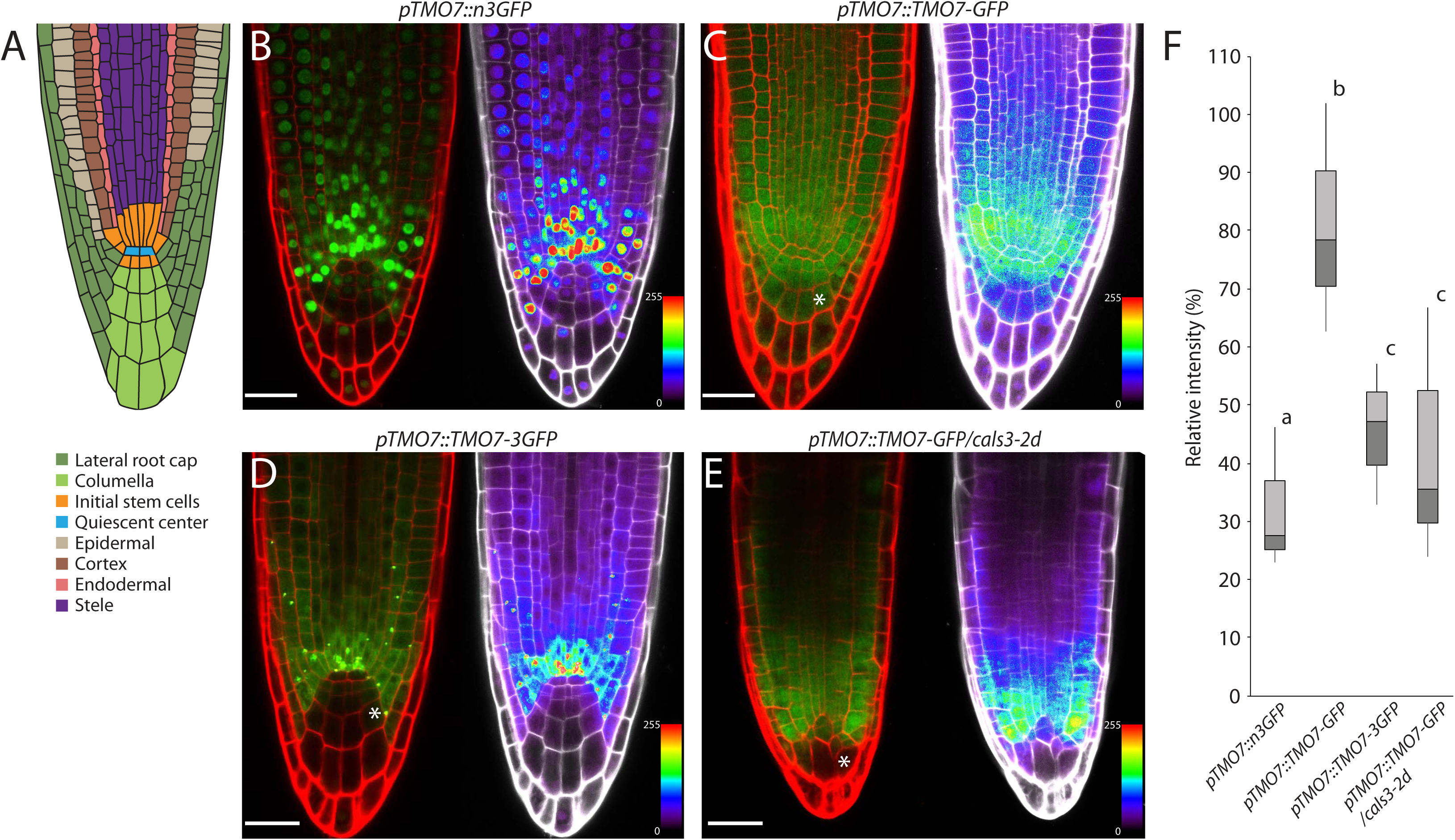
TMO7 moves through plasmodesmata into root cap cells in the Arabidopsis primary root. (A) Schematic image of Arabidopsis root structure. Different tissues are colour coded according to the colour legend. (B-E) 5-day-old root Fluorescence images in median plane of p*TMO7*::n3GFP (B), p*TMO7*::TMO7-GFP (C), p*TMO7*::TMO7-3GFP (D) and p*TMO7*::TMO7-GFP in *cals3-2d* (E); right panels show the false colour image of the left panel. (F) Statistical analysis of the fluorescence ratio between columella cells and stem cell niches regions of different transgenic plants, n≥13 roots per genotype, sample size is indicated in materials and methods. Significant differences by one-way ANOVA test are indicated by letters above bars (p<0.05). * marks the regions that shows highest movement differences. Scale bar = 25μm.

The finding that fusion to 3GFP prohibited transport of TMO7 suggests that TMO7 might, like most other mobile TFs, migrate through plasmodesmata which have a size restriction (Otero et al., 2016). To further investigate the possible passage of TMO7, we observed the movement of TMO7-GFP in the *cals3-2d* mutant, which over-accumulates callose and prohibits transport via PD (Vaten et al., 2011). Consistent with the notion that TMO7 moves through PD, the movement of TMO7-GFP was hampered in the primary root of *cals3-2d* mutants (Fig. 1E, F). Based on these observations we conclude that TMO7 protein moves through PD in the Arabidopsis root.

### TMO7 protein sequence, not size, instructs the unidirectional movement into root cap cells

Small proteins (molecule weight lower than 50 KDa) may transport freely through PD by diffusion (Crawford and Zambryski, 2001; Oparka et al., 1999). Since TMO7 is 93 amino acids long (around 11 KDa), it might thus passively diffuse through PD. To investigate whether movement is correlated to molecular weight, or rather a consequence of specific protein features, we focused on other small bHLH proteins. The Arabidopsis genome encodes four TMO7-like proteins (T7L1-4; De Rybel et al., 2011), all representing small bHLH TFs (92 to 94 a.a.; Fig. 2A, B). Within the *TMO7* clade, amino acid sequence is very conserved (up to 45% identity, and 80% similarity; Fig. 2B). We therefore also selected two other small bHLHs outside the *TMO7* clade, *AtbHLH138* (AT2G31215, 129 a.a., about 15 KDa) and *AtbHLH151* (AT2G47270, 102 a.a., about 12 KDa) (Fig. 2A, B), to uncouple size from homology. All small bHLH proteins were expressed as GFP fusion proteins from the *TMO7* promoter, and we visualized localization (Figure 2C, D) and quantified mobility (Figure 2E).

**Fig 2.**
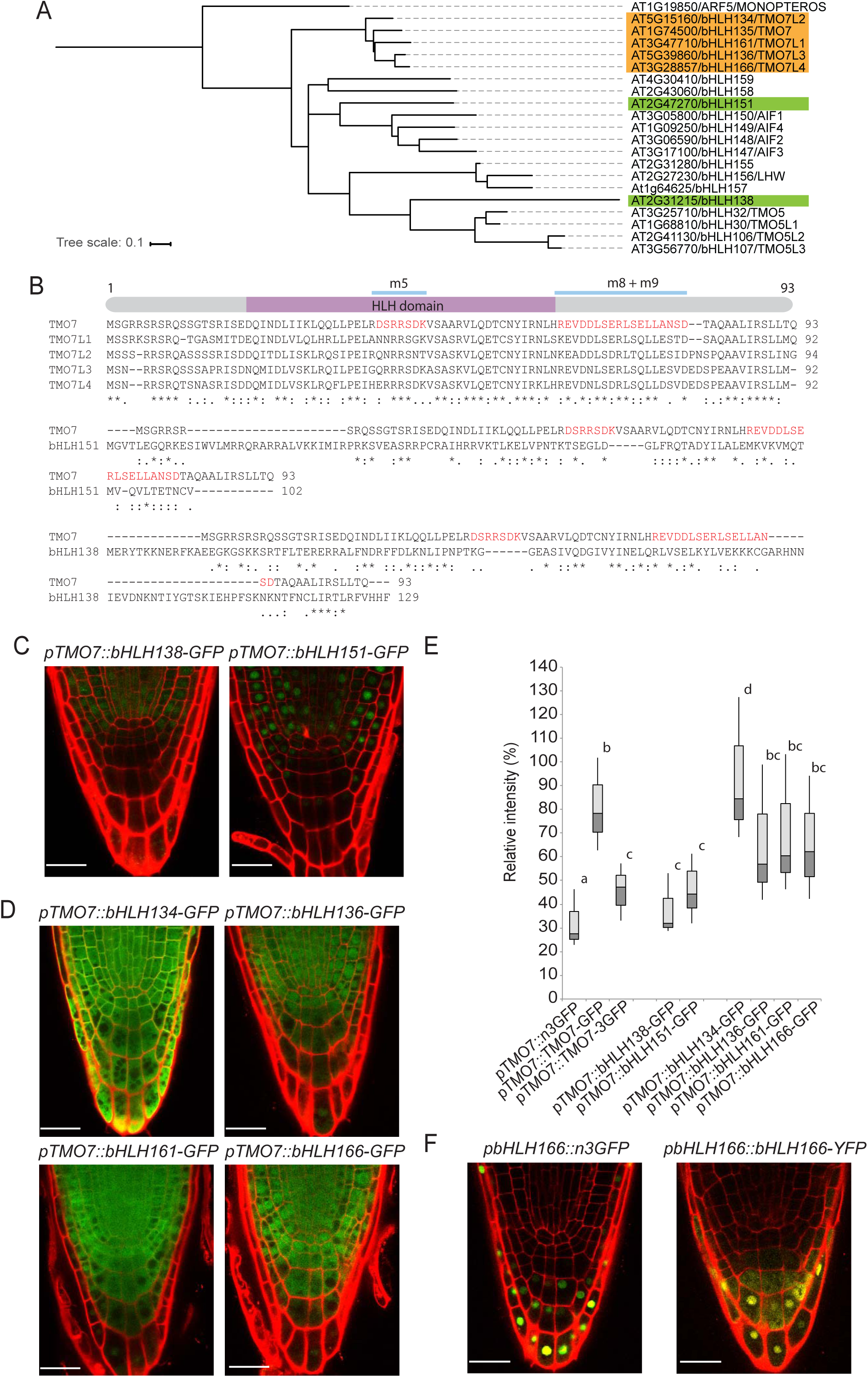
TMO7 family proteins carry intrinsic features required for transport. (A) Phylogenetic analysis of selected bHLH proteins in Arabidopsis based on the full-length protein sequence. Branch lengths indicate phylogenetic distances (see scale bar: fraction of deviations). Green rectangle indicates the location of bHLH138 and bHLH151. ARF5/MP is used as an unrelated reference. (B) Protein sequences comparison within the TMO7 family and between TMO7, bHLH138, and bHLH151. The schematic figure indicates the location of HLH domain and the mobile *cis*-elements in TMO7 protein in red. (C-E) Mobility analysis of bHLH138, 151 and TMO7-Like proteins. Confocal images of bHLH138-GFP, bHLH151-GFP (C), and bHLH134-, bHLH136-, bHLH161-, and bHLH166-GFP expressed by the *TMO7* promoter (D), and the fluorescence ratio analysis (E) Samples size is indicated in materials and methods. Significant differences by one-way ANOVA test are indicated by letters above bars (p<0.05). (F) Confocal images of p*bHLH166*::3nGFP and p*bHLH166*::bHLH166-GFP, note that no fluorescence was detected outside the promoter active regions. Scale bar = 25μm.

We observed comparable mobility of all proteins within the TMO7-like clade (Fig. 2D, E). Interestingly, unlike other T7Ls, the bHLH134-GFP protein localized not only in the nucleus and cytoplasm but also strongly at the cell periphery (Figure 2D). Nonetheless, mobility of bHLH134-GFP was not influenced by this altered localization (Fig. 2D, E). The unrelated bHLH proteins, bHLH138 and bHLH151, localize mainly to the nucleus (Figure 2C), consistent with the presence of several basic residues (K, R – basic residues constitute nuclear localization signals) N-terminal to the HLH domain (Figure 2B). In addition, bHLH138 also partially localized in the cytoplasm (Fig. 2C). Yet, despite their small size, no movement of bHLH138 or bHLH151-GFP could be detected (Fig. 2C, E). Thus, we conclude that small protein size does not determine TMO7 movement. Rather, the absence of a clear nuclear localization signal and/or the presence of conserved motifs within the TMO7 clade proteins may mediate transport through PD.

We further explored the transport at the root columella cells. We previously reported promoter activity of *T7Ls: bHLH134, bHLH136*, and *bHLH166* are expressed weakly in the root cap and columella cells while *bHLH161* is expressed only in the lateral root cap (De Rybel et al., 2011). We therefore generated YFP fusion protein with the endogenous promoter to explore the direction of transport. Among these fusion proteins, expression could only be detected for bHLH134-YFP (not shown) and bHLH166-YFP (Fig 2F). Interestingly, both fusion proteins showed highest intensity at the very tip of the columella cells with decreasing intensity toward the shoot, which is comparable to their promoter activity (Fig 2F; De Rybel et al., 2011). Thus, we conclude that while bHLH134 and 166 proteins are rootward mobile when expressed in the TMO7 expression domain, they do not normally move shootward from their expression site. This finding also suggests that the transport of TMO7-like proteins is unidirectional between meristematic cells and root cap cells.

### Protein sequence elements and subcellular localization define TMO7 transport

Given that the TMO7 protein sequence directs transport, we aimed to map the crucial region(s) by systematic mutation. We hypothesized that mutations which affect TMO7 mobility elements would disrupt transport. We performed a linker-scanning analysis by replacing regions of seven to nine amino acids with an poly-Alanine linker of the same length, and thus generated eleven TMO7 mutants (p*TMO7::*TMO7_*m1*_-GFP to p*TMO7::*TMO7_*m11*_-GFP; Fig. 3A). The relative intensity of fluorescence in the tip versus meristem was quantified in more than 10 independent transgenic lines for each mutant (except for *m10;* only 5 T2 lines), and lines showed some variability in transport (Figure 3C). Yet, transport appeared unaffected in most mutant versions (Figure 3B, C). We found consistent reduction in movement for three mutant proteins, TMO7*_m5_*, TMO7_*m8*_ and TMO7_*m9*_, similar to the pattern found in *pTMO7::*TMO7-3GFP (Fig. 3B, C). In addition, TMO7_*m2*_- and TMO7_*m3-*_GFP showed movement defects in some, but not all transgenic lines (Fig. 3B, C). These results demonstrate that movement of TMO7 depends on at least two major and one minor elements.

**Fig 3.**
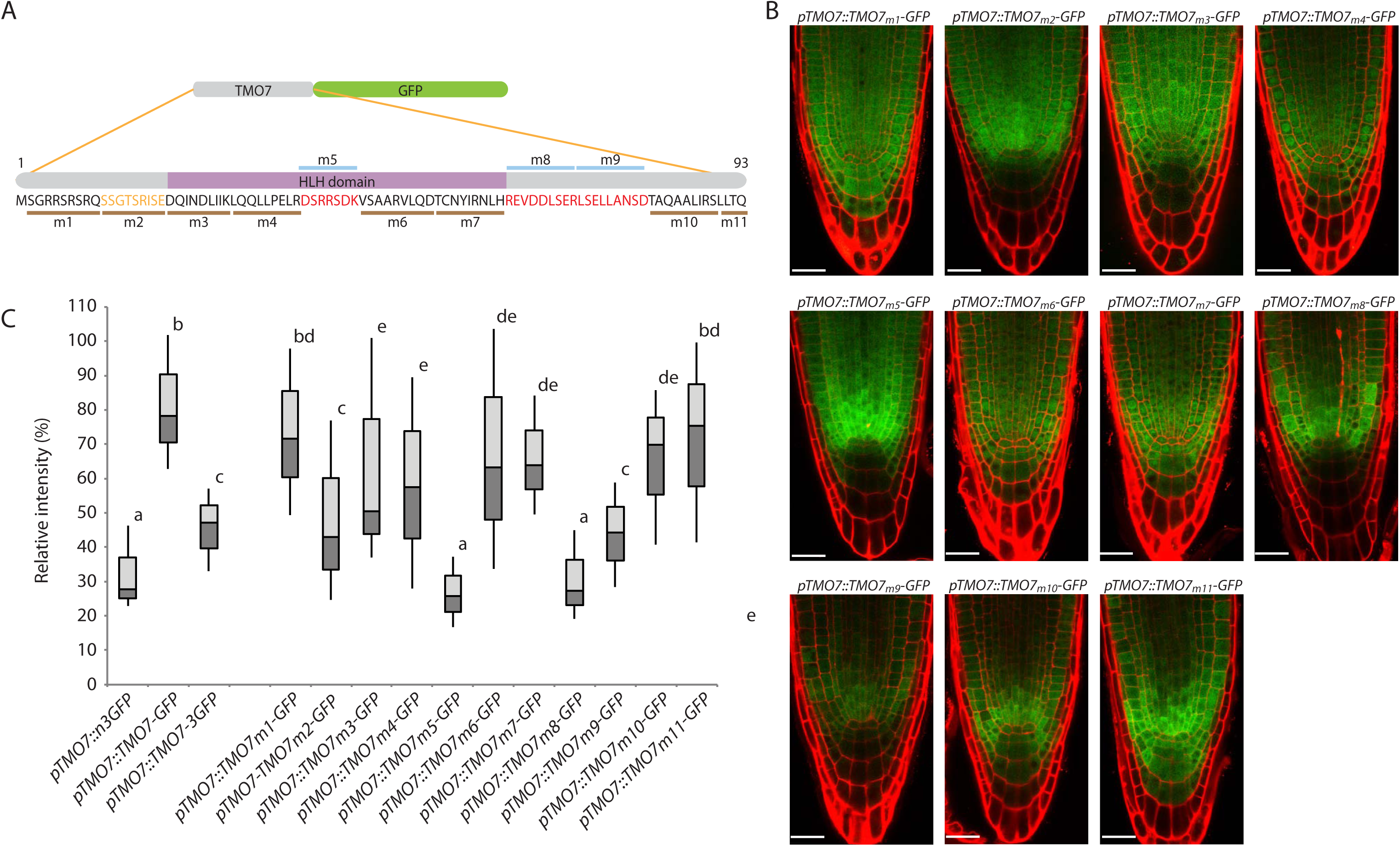
TMO7 contains two major and one minor mobility motifs. (A) Schematic figure indicating the position of each mutant, those indicated above the sequence are mobile defective mutants. (B) Confocal images of each linker-scanning mutant. Note that the QC and columella region in m2, m5, m8 and m9 region shows reduced or no fluorescence. (C) Fluorescence ratio analysis of linker-scanning mutants, sample size is indicated in materials and methods. Significant differences by one-way ANOVA test are indicated by letters above bars (p<0.05). Scale bar = 25μm.

By comparing the sequence between TMO7-like family proteins and the immobile bHLH138/151 proteins, we found the M5 region to locate within the conserved HLH domain and with very high conservation between all TMO7 families. However, bHLH138 and bHLH151 have limited similarity with the M5 region (Fig 2B). The M8/9 regions locate immediately C-terminal to the HLH domain, and this region is also highly conserved within the TMO7 family, but different from bHLH138 and bHLH151 (Fig 2B). In addition, we also compared the bHLH domain of GLABRA3 (bHLH1), a known mobile protein that regulates root hair formation (Bernhardt et al., 2005), with TMO7 and we only found limited similarity at the M5 or M8, 9 regions (data not shown), indicating that tissue-specific transport systems may operate in the Arabidopsis root. These data suggest that M5 and M8/9 regions may be specific elements responsible for mobility of TMO7 family proteins.

An intuitive hypothesis would be that the *m5* and *m8/9* mutations alter the secondary or tertiary structure of the TMO7 protein and thus lead to movement defects. We analysed the possible secondary structure with the SWISS-MODEL protein-folding tool (http://swissmodel.expasy.org/interactive). The predicted TMO7 structure is very similar to the bHLH transcription factor MyoD (Fig S2B; Ma et al., 1994); however, none of the mutants that disrupt movement is predicted to have a different secondary structure (Fig. S2A). It is therefore not directly evident how the mutations would affect TMO7 protein properties, but given that no alternate secondary structures are predicted, the mutations may also not have dramatic consequences for protein folding.

To further test whether M5, M8 and M9 are *bona fide* mobility elements, sufficient for driving movement, we inserted these elements into bHLH138 and bHLH151 to create chimaeric proteins. Since the syntax surrounding M5 and M8/9 may also influence their function, chimaeric proteins were designed based on the syntax of TMO7. Based on the protein-folding prediction, M5 is localized at the loop region of HLH domain (Fig. S2B). We replaced the loop sequence of bHLH138 and 151 by the TMO7 M5 sequence to create bHLH138-M5 and bHLH151-M5. Since there is no predicted secondary structure at the M8/9 region, and because it directly follows the HLH region in TMO7, we directly inserted the M8/9 region after the bHLH domain of bHLH138 and 151 to generate bHLH138-M8/9 and bHLH151-M8/9 respectively. The combination of M5 and M8/9 was also constructed as bHLH138-M5/8/9 and bHLH151-M5/8/9 (Fig. 4). All the chimeric bHLH138 and bHLH151 versions except bHLH151-M5-GFP slightly gained movement (Fig. 4B). However, compared to TMO7, movement of chimaeric bHLH138/151 was limited (Fig. 4B), perhaps due to strong nuclear localization of bHLH138 and 151 (Fig. 2C and 4A). It was previously shown that movement of mobile proteins can be altered by localizing the protein to specific subcellular compartments (Crawford and Zambryski, 2000; Daum et al., 2014; Gallagher et al., 2004; Kim et al., 2005; Rodriguez et al., 2016; Tamaki et al., 2007).

**Fig 4.**
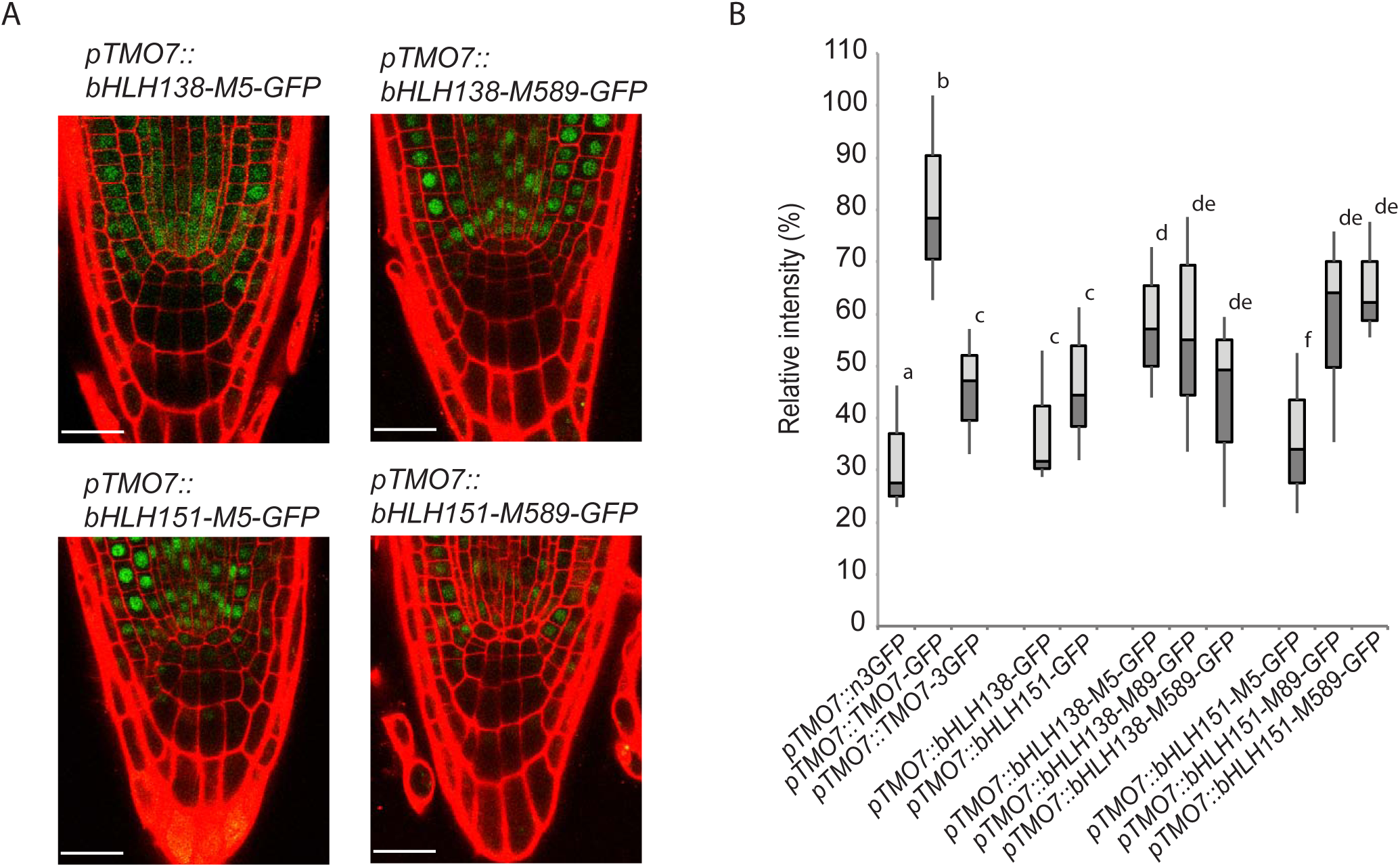
Transferability of TMO7 mobility motifs. (A) Confocal images of bHLH138, bHLH151 with M5 and M5, 8, 9 insertions. (B) Fluorescence intensity analysis of bHLH138-M5, -M8,9, -M5, 8, 9-GFP and bHLH151-M5, -M8,9, -M5, 8, 9-GFP. Sample size is indicated in materials and methods. Significant differences by one-way ANOVA test are indicated by letters above bars (p<0.05). Note that the images have been enhanced for visualization. Scale bar = 25μm.

However, in contrast to a sole movement-restricting capacity of nuclear localization, in our linker scanning analysis, we also often observed a lack of nuclear protein in movement-defective mutants (Fig. 3B and 5A). This suggests that transport into the nucleus may contribute to TMO7 transport. To test whether strong subcellular localization interferes with the mobility of the protein, we generated TMO7-GFP protein with SV40 nucleus localization (NLS) or nuclear export signals (NES) at the N- or C-terminus of TMO7-GFP, respectively (NLS-TMO7-GFP; TMO7-GFP-NES). Like in other mobile proteins (Balkunde et al., 2017; Gallagher et al., 2004; Rodriguez et al., 2016), the nuclear localization signal restricted TMO7 movement. Surprisingly however, the NES also hindered its mobility (Fig 5B-D), suggesting that entering as well as leaving the nucleus might be crucial for TMO7 transport.

**Fig 5.**
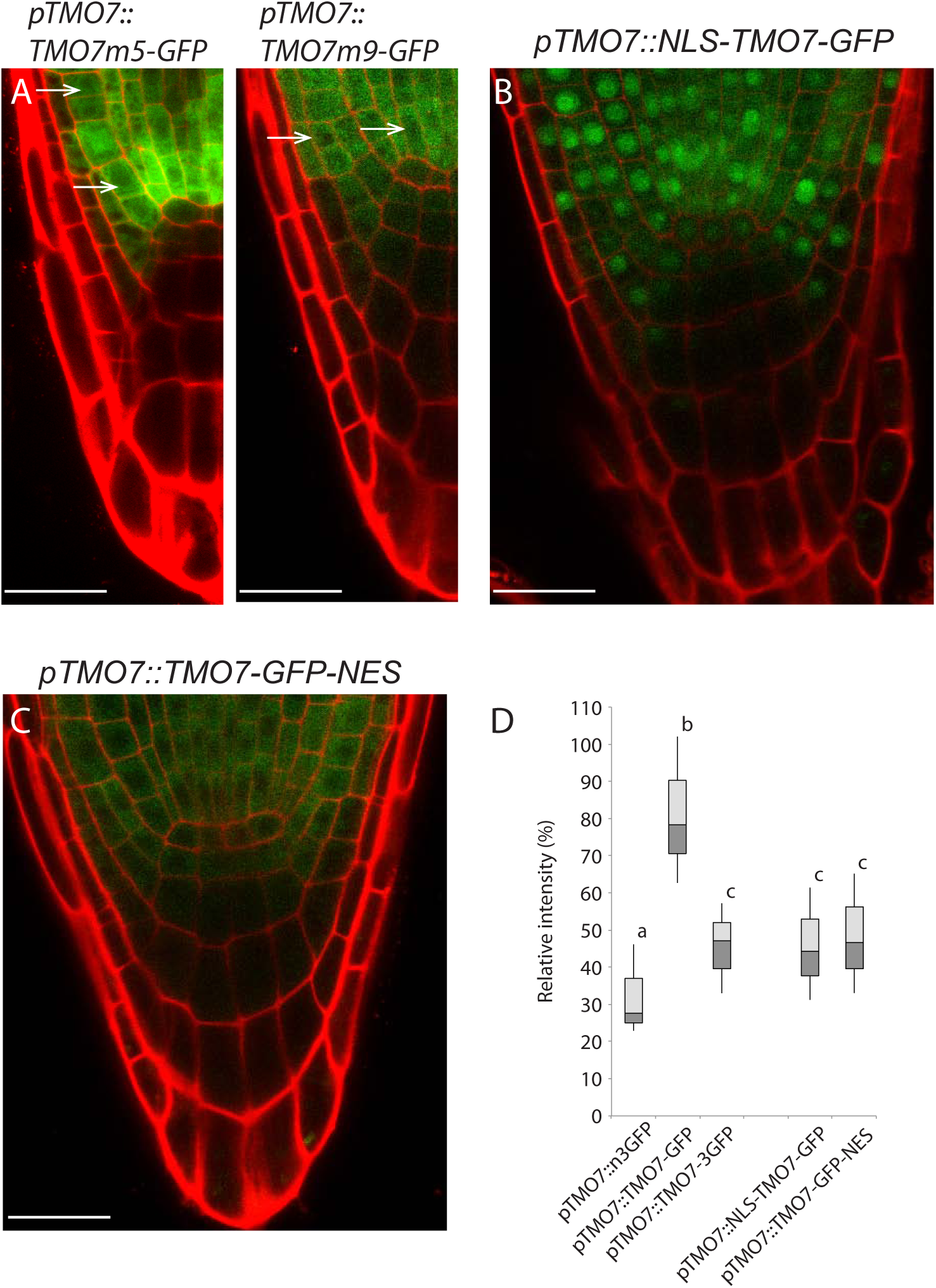
Nuclear localization and exclusion signals restrict the movement of TMO7. (A) Confocal image of the p*TMO7::TMO7m5*-GFP and p*TMO7::TMO7m9*-GFP root tip. Arrows indicate nucleus without fluorescence. (B and C) Confocal images of p*TMO7*::NLS-TMO7-GFP (B) and p*TMO7*::TMO7-GFP-NES (C), note that fluorescence signal is absent in the root tip regions in both images. (D) Fluorescence intensity analysis of p*TMO7*::NLS-TMO7-GFP and p*TMO7*::TMO7-GFP-NES, sample size is indicated in materials and methods. Significant differences by one-way ANOVA test are indicated by letters above bars (p<0.05). Scale bar = 25μm.

In summary, the TMO7 family contains two major mobile *cis*-elements which partially endow mobility of bHLH138/151 when transplanted. However, the mobility strongly depends on subcellular localization; the strong NLS decreases the potential of direct interaction between TMO7 and PD, while the result of the TMO7-GFP-NES suggests that the import of TMO7 into nucleus is also critical for its movement.

### Phosphorylation may control TMO7 mobility

Given that specific motifs, as well as residence in the nucleus, seem important for TMO7 movement, we hypothesized that TMO7 transport involves post-translational modification. To start exploring this option, we first analysed the potential of phosphorylation on TMO7 using a prediction server (http://www.dabi.temple.edu/disphos/). Among the 19 Serine, Threonine and Tyrosine residues, phosphorylation was predicted to occur on 8 (Table S2). Among these, S39 and S42 are located within the M5 region (Fig. 3A). We therefore substituted each and both of the Serine residues by Alanines to create p*TMO7::*TMO7-S39A-, S42A- and S39, 42A-GFP, respectively. The subcellular localization of TMO7 was not altered by the mutations, indicating that protein import into the nucleus is not impaired by the mutation (Fig. 6A, B and C). Interestingly however, all mutants showed impaired mobility (Fig. 6A-D), which suggests that both of these potential phosphorylation sites are important for TMO7 transport.

**Fig 6.**
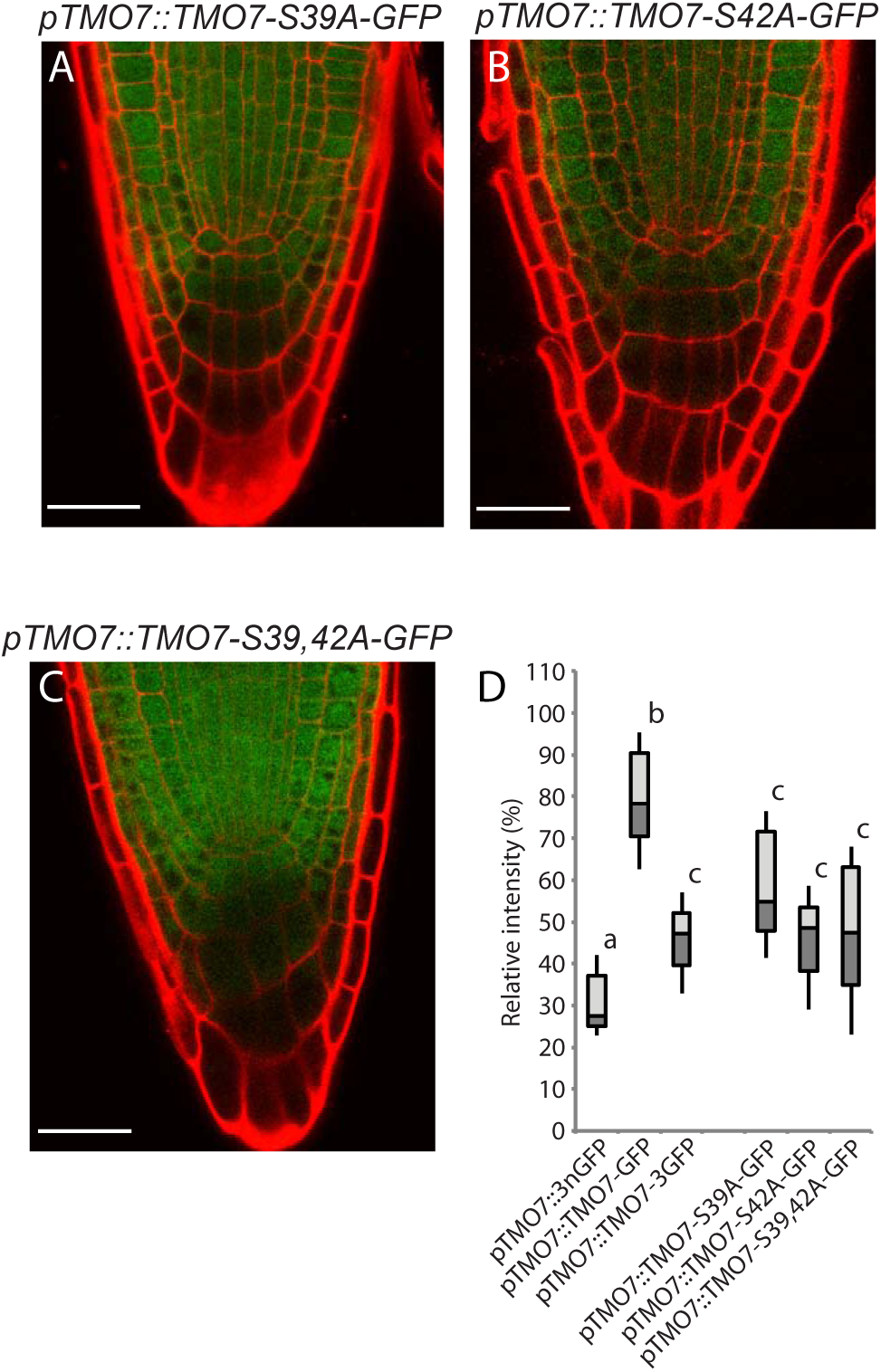
Potential phosphorylation sites contribute to TMO7 movement. (A-C) Confocal images of p*TMO7::*TMO7S39A-, S42A-, and S39, 42A-GFP. (D) Fluorescence intensity analysis of p*TMO7*::TMO7S39A-, S42A-, and S39, 42A-GFP, sample size is indicated in materials and methods. Significant differences by one-way ANOVA test are indicated by letters above bars (p<0.05). Note that the expression intensity of the TMO7 mutants is relatively lower compared to the p*TMO7*::TMO7-GFP line, however, the statistical analysis still indicates the reduction of mobility in all the mutants. Note that the images were modified for visualization. Significant differences by one-way ANOVA test are indicated by letters above bars (p<0.05). Scale bar = 25μm.

### Characterization of a stable *tmo7* mutant reveals complex regulatory interactions

To address the importance of TMO7 transport, we first aimed to generate lost-of-function resources. Previously, we described a mild hypophysis division phenotype, as well as a low-penetrance rootless seedling defect in lines that had the *TMO7* gene silenced using RNAi or artificial microRNA expression. No phenotype could be observed in the available *tmo7-1* and *tmo7-2* insertion mutants (Schlereth et al., 2010). It is possible that RNAi and amiRNA approaches targeted homologs as well as *TMO7*, or alternatively it could be that the *tmo7-1* and *tmo7-2* insertion lines may not represent null alleles. Thus, we generated mutants through CRISPR-Cas9 gene editing (Tsutsui and Higashiyama, 2016). We designed a short-guide RNA targeting the M5 mobile element and obtained 3 independent mutant alleles (*tmo7-4, tmo7-5* and *tmo7-6*), each creating shorter and mutated predicted proteins (Fig. 7A, B). In all CRISPR *tmo7* alleles, we observed that root length was reduced compare to wild-type plants (Figure 7C, D). In addition, we consistently found hypophysis division defects in all three mutant alleles (*tmo7-4* = 12.4%, n= 331; *tmo7-5* = 5.6%, n=302; *tmo7-6* = 5%, n=357; Col = 1.8%, n= 277; Fig 7E). However, the rootless phenotype that was observed in 1-7% of seedlings in *amiRTMO7* and *TMO7 RNAi* lines (Schlereth et al., 2010), was not recovered in the *tmo7-4, tmo7-5* and *tmo7-6* alleles (n>1000 seedlings for *tmo7-4* and *tmo7-5*, n>600 for *tmo7-6*). This suggests that the hypophysis defect might later be repaired by a yet unknown mechanism in *tmo7-4, -5*, and *-6* mutants.

**Fig 7.**
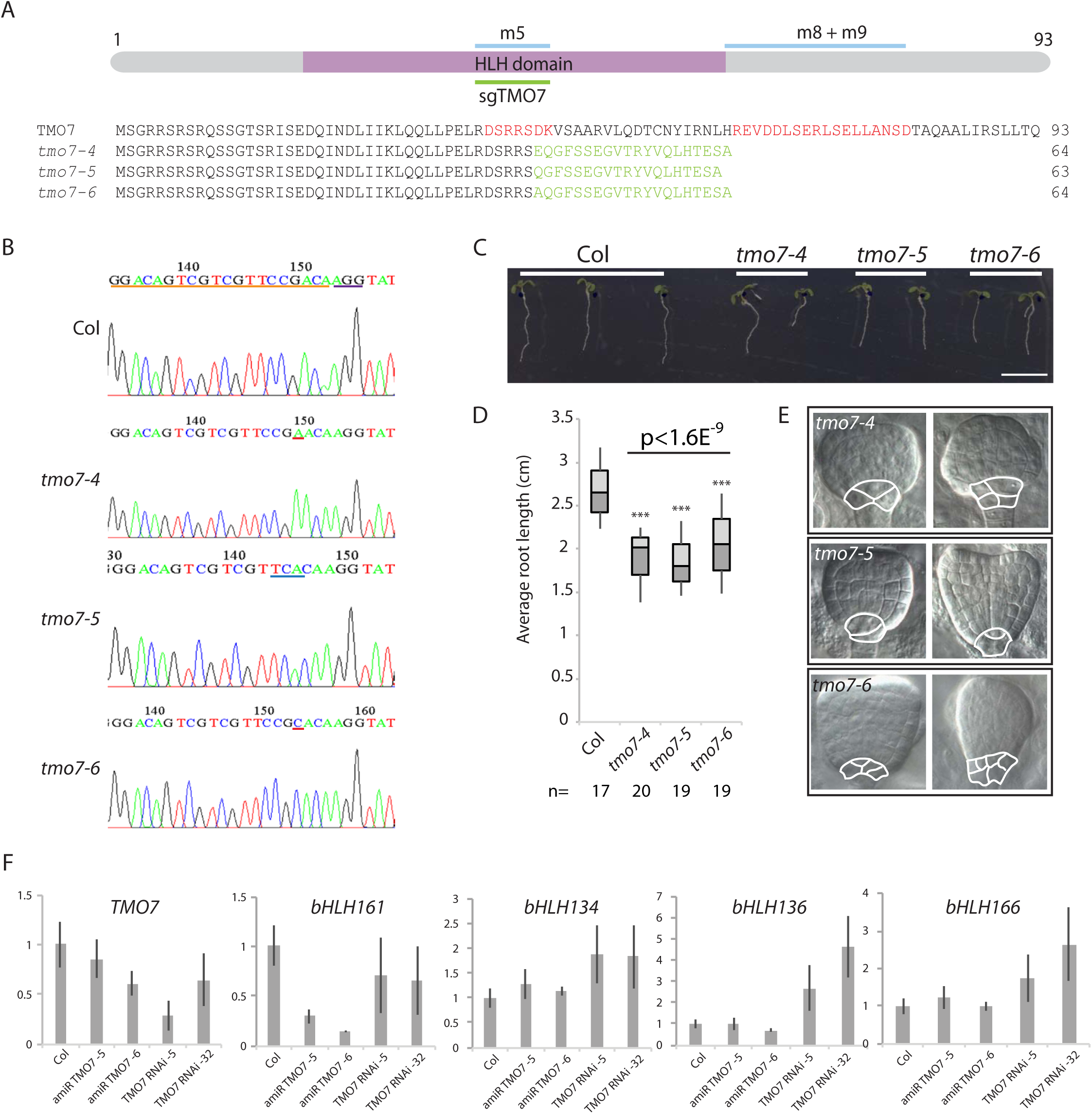
Characterization of CRISPR-Cas9 *tmo7* mutants. (A) Schematic figure indicates the sgRNA targeting region and the mutated amino acid sequences (in green) of *tmo7-4, tmo7-5*, and *tmo7-6*. (B) Sequencing traces of Col, *tmo7-4, tmo7-5* and *tmo7-6*. Orange line indicates sgRNA target sequence, purple line indicates the protospacer adjacent motif (PAM) in Col. The red line indicates the inserted nucleotide in *tmo7-4* and *tmo7-6*, the blue line indicates the deletion region in *tmo7-5*. (C) Root structure of Col, *tmo7-4, tmo7-5* and *tmo7-6*. Scale bar= 1cm. (D) Average root length comparison between Col and CRISPR *tmo7* lines, significance by student t-test, ***p<0.01. (E) Differential interference contrast (DIC) images of hypophysis division phenotype observed in *tmo7-4, tmo7-5*, and *tmo7-6*. Scale bar= 1cm. (F) Expression level of *TMO7* family genes in RNA suppression lines by qRT-PCR analysis, relative expression level compared with endogenous control, *ACTIN2*. Quantified with three biological repeats and error bar indicates standard error of mean.

One possibility is that the TMO7-like proteins act redundantly with TMO7; alternatively and additionally, *TMO7-like* genes may be involved in later stages and help generate an embryonic root following initial defects upon TMO7 depletion. In either scenario, it would be expected that the expression of *TMO7-like* genes is misregulated upon *TMO7* gene silencing, for example as an off-target effect. To test this possibility, we quantified *TMO7Ls* expression in primary roots of TMO7 RNA suppression lines by quantitative reverse transcription poly-chain reaction (qRT-PCR). The results confirm that the *TMO7* transcript is down-regulated between 15 and 70% in both *amiRTMO7* and *TMO7 RNAi* lines (Fig. 7F). Furthermore, we observed complex changes in the regulation of *TMO7* homologues, as different *TMO7-like* genes are down-regulated in different gene silencing lines, while others are up-regulated (Fig. 7F). In all *tmo7* CRISPR alleles, *TMO7* and all *TMO7-like* genes were expressed at a level comparative to wild-type (data not shown). These results are consistent with the notion that redundancy among *TMO7-like* genes may contribute to embryonic root formation. In RNA suppression lines, multiple homologs are affected, which may be the cause for the rootless phenotype. Clearly, a higher-order (CRISPR/Cas9) mutant, knocking out *TMO7-like* genes as well as *TMO7*, will be required to explore the function of *TMO7* in embryonic root formation.

### TMO7 movement contributes to hypophysis division

Likely functional redundancy with close relatives prevents us from directly testing whether TMO7 movement is critical for hypophysis and embryonic root development. However, we employed the *cals3-2d* mutant to indirectly address this question. The hyper-accumulation of callose in *cals3-2d* mutant prohibits the movement of proteins, and we noticed that the mutant (Vaten et al., 2011) shows a strong rootless phenotype, resembling the *monopteros* (*mp*) (Berleth and Jurgens, 1993) and *TMO7 RNAi/amiRTMO7* phenotype (Fig 8A). A detailed analysis of cell division patterns in the embryo showed a lack of hypophysis division in nearly all *cals3-2d* mutant embryos (Fig 8C, D; 95%; n>700 embryos). As plasmodesmata were reported to be involved in auxin transport (Han et al., 2014), and because auxin transport from embryo to hypophysis promotes root formation (Friml et al., 2003; Weijers et al., 2006), we first tested if the hypophysis and root defect in the *cals3-2d* mutant is related to an auxin accumulation problem. We transformed an auxin response reporter, p*DR5::*nGFP, (Weijers et al., 2006) into the *cals3-2d* background. Strikingly, the expression of p*DR5*::nGFP in the hypophysis was not affected by *cals3-2d* (Fig 8B), which suggests that auxin is normally transported to the hypophysis in the mutant.

**Fig 8.**
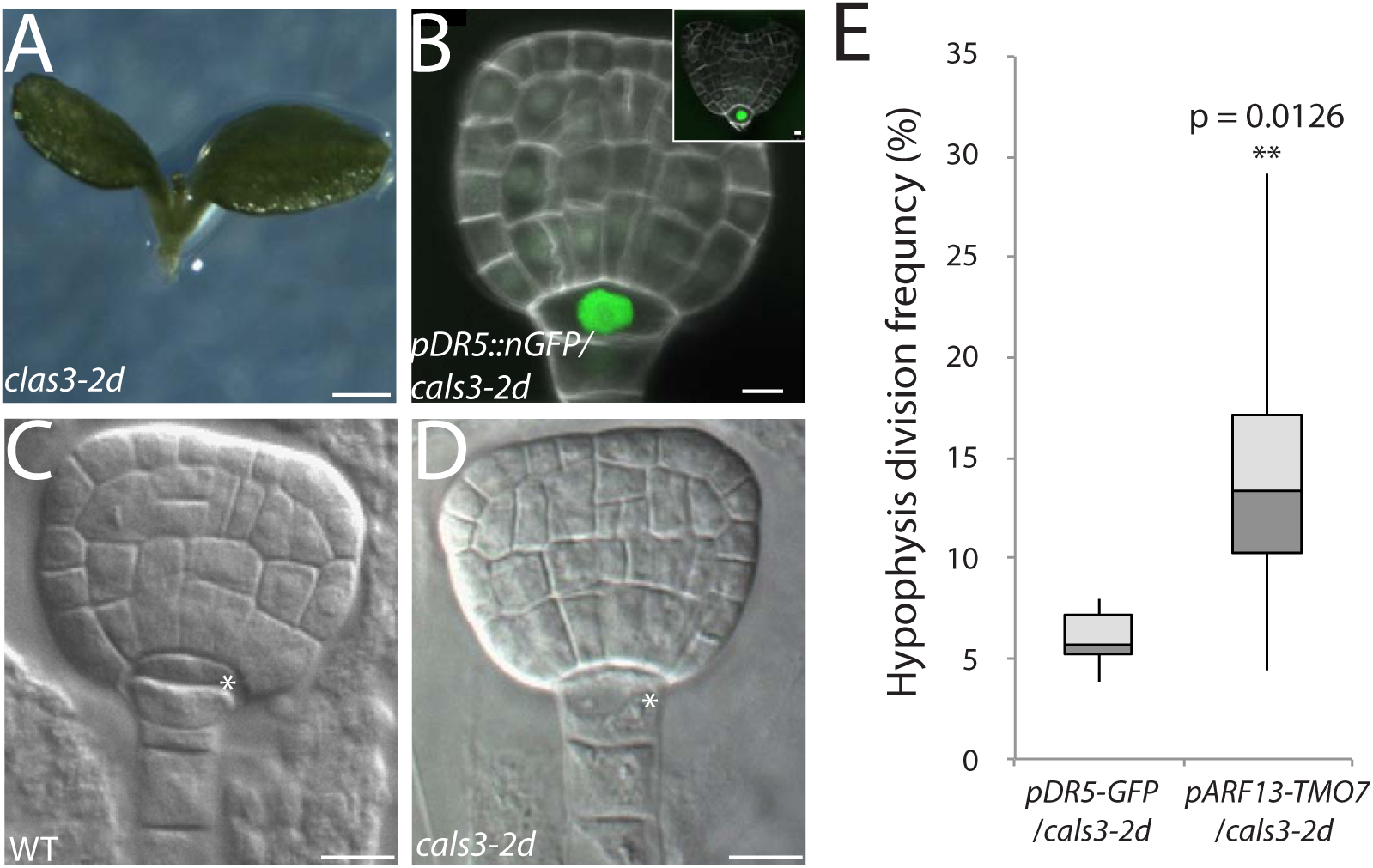
Reintroduction of TMO7 into the *cals3-2d* suspensor partially complements the hypophysis division phenotype. (A) Rootless phenotype of *cals3-2d* mutant. Scale bar = 100μm. (B) Expression of p*DR5::*nGFP in a *cals3-2d* embryo. Note that the accumulation of GFP signal is in the upper-most suspensor cell, and still no division has occurred in heart stage (insert). Scale bar = 5μm. (C and D) DIC image of a wild-type early-globular stage embryo (C) and a *Cals3-2d* late-globular stage embryo (D). Scale bar = 10μm. Note that the division that normally occurs in WT (asterisk mark in C) is absent in *cals3-2d* embryo (asterisk mark in D). (E) Statistical analysis of wild-type division frequency in p*DR5*::nGFP/*cals3-2d* (n=7 independent lines, at least 100 embryos per line)and p*TMO7*::TMO7-GFP/*cals3-2d* (n=9 independent lines, at least 90 embryos per line) embryos. **: 0.01<p<0.05, student t-test.

Due to difficulties with detecting the low TMO7 protein amounts in the embryo, we could not determine if TMO7 movement is affected in the *cals3-2d* mutant embryo (not shown). However, given that TMO7 movement in the root of the *cals3-2d* mutant is impaired (Fig 1E, F), we tested if expression of TMO7 in the cells to which it is normally transported (uppermost suspensor cells) would alleviate the division defect in the *cals3-2d* mutant. Therefore, TMO7 was expressed from the suspensor-specific *ARF13* promoter (Rademacher et al., 2012). By counting the correct division frequency of the hypophysis in both p*ARF13::*TMO7 and the p*DR5::*nGFP control, we found a partial rescue of hypophysis division in *cals3-2d* by p*ARF13*::TMO7 expression (Fig 8E; 14.4 ± 7.3% with p*ARF13*::TMO7 compared to 6.0 ± 1.5% with p*DR5*::nGFP; n=7 and 9 lines respectively with at least 90 individuals for each line; p=0.0126 by two-tail student t-test). These results suggest that the hypophysis division phenotype in the *cals3-2d* mutant is likely caused by PD obstruction, which prevents TMO7 movement, but not auxin transport. Our findings thus indicate that TMO7 mobility contributes to hypophysis division.

## Discussion

In this study, we systematically analysed the mobility of a specific small bHLH transcription factor family protein. We conclude that transport of TMO7 is sequence-dependent, but not primarily determined by protein size. Interestingly, the shuttling into and out of the nucleus appears to be critical for mobility (Fig. 5). We postulate that this shuttling into the nucleus labels the protein for transport to other cells. The importance of nuclear localization for a mobile protein has also been proposed for the transport of SHR and another mobile R3-type MYB-like protein, CAPRICE (CPC), which expresses in the non-hair epidermal cells in the roots and moves to the hair cells to specify the formation of the root hairs (Gallagher and Benfey, 2009; Gallagher et al., 2004; Kurata et al., 2005). A T289>I point mutation in SHR, prevents the import of SHR into the nucleus and its intercellular transport while the W76A mutation of CPC largely reduced its nuclear localization and prevented its movement (Gallagher et al., 2004; Kurata et al., 2005). The Tyrosine residue 289 of SHR was predicted to be a phosphor-acceptor which functions in dimerization and proper localization (Gallagher et al., 2004). Consistent with the role of T289 in SHR, the serine residues (S39 and S42) in TMO7 were also predicted to be phosphorylated. Replacing the residues by Alanine hampers the intercellular movement, which suggests that phosphorylation might be a mark for intercellular transport (Fig 6). However, we cannot rule out the possibility that these mutations act independently of phosphorylation by e.g. distorting the protein structure or preventing the interaction with an unknown factor which might be critical for transport, but not for nuclear import. Further analysis of the post-translational modification status of mobile proteins will likely provide more insight to intercellular communication mechanisms.

We previously showed that TMO7 is only transported into the hypophysis, but not into the upper tier of the pro-embryo (Schlereth et al., 2010). Consistent with this directional transport property in the embryo, we found that TMO7 family proteins can move from the root meristematic region into the root cap, but not in the opposite direction (Fig 2D, F). In addition, in our p*TMO7*::TMO7-3GFP line, we observed obvious aggregations in the seedling root which is lacking in p*TMO7*::TMO7-GFP and other TMO7Ls-GFP fusion lines (Fig 1 and 2). The aggregations seem to be highly enriched in the meristematic cells closest to the neighbouring QC (Fig. 1D). It is possible that these clusters of proteins are TMO7 with intercellular transport markers but over the limit size for passing through PD. Interestingly, the stem cell maintenance transcription factor WUSCHEL HOMEOBOX 5 (WOX5) has also been shown to move from the QC into the columella stem cells, and no shootward transport was observed (Pi et al., 2015). These data suggest that different regulatory mechanisms exist to control the rootward and shootward transport between root cap cells and other meristematic niche cells.

The cell walls between the pro-embryo and suspensor and between the upper and lower tiers in the embryo are generated by the very first and second cell divisions during embryogenesis (Scheres et al., 1994; ten Hove et al., 2015). The symplastic transport between suspensor and pro-embryo and within the pro-embryo was shown to be liberal until the globular stage, as soluble GFP moved from the suspensor to the whole embryo when expressed from the *AtSUC3* promoter and across the whole embryo when expressed from the *AtSTM* promoter (Stadler et al., 2005). Interestingly, the shootward mobility of TMO7 to upper-tier cells is limited in the early embryo, which suggests the symplastic transport control between the two tiers is established early in embryogenesis. How and why this control is established, and at which developmental stage is a very intriguing question.

In *tmo7-4, -5*, and *-6*, we observed a clear hypophysis phenotype, which is consistent with the previous finding that *TMO7* is involved in hypophysis development. Interestingly however, no rootless phenotype was recovered. A possible scenario is that in the early embryo, TMO7 together with other mobile factors coordinate the development of hypophysis while other TMO7 family genes control the later development of QC and root cap. This scenario could explain why, hypophysis errors in *tmo7* CRISPR mutants do not lead to rootless defects. Indeed, both *TMO7* gene silencing lines and *tmo7* CRISPR lines give rise to hypophysis defects but only the gene silencing lines show a low-frequency rootless phenotype, in which *TMO7* family genes were misregulated (Fig 7F). It has been shown that the *TMO7* family genes are not expressed at least until the late heart stage during embryogenesis but are expressed in the post-embryonic root in the columella and lateral root cap (De Rybel et al., 2011). Therefore, the misregulation of *TMO7* family genes in RNA suppression lines might partially impair a redundancy-based repair mechanism. A higher-order *TMO7* family mutant might reveal how *TMO7* family genes help the development of hypophysis and post-embryonic root, and may also provide insight into the genes and cellular pathways that are controlled by mobile TMO7/T7L proteins in root formation and development.

## Materials and methods

### Plant materials

All seeds were surface sterilized, sown on 1/2 strength MS with 0.01% MES, 1% sucrose and 0.8% Daishin agar plates, and stratified for 1 day at 4°C in the dark before they were grown under long-day (16/8 hr) conditions with a constant temperature of 22°C in a growth room. The TMO7 reporter lines, p*TMO7::*3nGFP, p*TMO7::*TMO7-GFP and p*TMO7::*TMO7-3GFP have been previously described (Schlereth et al., 2010). The *cals3-2d* mutant (Vaten et al., 2011) was kindly provided by Ykä Helariutta (SLCU, Cambridge).

### Cloning

All constructs except those for CRISRP-Cas9 gene editing were generated by LIC cloning methods with pPLV02 vectors as previously described (De Rybel et al., 2011). For mobility analysis of bHLH138/151, TMO7 family proteins, and TMO7 S39, 42A mutants, a vector containing *TMO7* promoter (*pTMO7* vector) was first created by amplification of a 2.2kb promotor fragment from *pTMO7-n3GFP* that was subsequently ligated through *EcoRI* and *BamHI* sites with the pPLV02 vector. The target genes and GFP were amplified separately with overlapping linkers and further fused by fusion PCR with primers containing the LIC adaptor. The TMO7 S39, 42A mutants were amplified using site direct mutagenesis primers. The 5’ and 3’ fragments were further fused together by PCR primers containing the LIC adaptor. The fused fragments were integrated with the *pTMO7* vector by LIC cloning. For Alanine-linker scanning, the 2.2kb promoter of *TMO7* plus the 5’ fragments before the mutation site and the 3’ fragments after the mutation site plus the GFP tag were separately amplified by PCR. The two fragments were fused by PCR with primers containing the LIC site and further integrated with pPLV02 by LIC cloning. All primers are listed in table S1.

### Microscopic and expression analysis

For imaging of roots, 5-d-old-seedlings were incubated in 10 mg/mL propidium iodide solution for 1-2 minutes. GFP and propidium iodide were visualized by excitation at 488 nm and detection between 500 to 535 nm and 630 to 700 nm, respectively. For florescence ratio analysis, the two regions of interest (ROI) were selected in the Leica Application Suite (LAS) program as in Fig S1. The QC and three layers of columella cells were selected as ROI1; the lateral root cap cells and cells in the stem cell niche (up to the twelfth cortex cell) were selected as ROI2. To prevent fluorescence intensity variation due to the tilting of the root, fluorescence of PI staining was used as the reference. The intensity ratio between the two channel of individual ROI was first calculated, and the final intensity ratio was calculated by dividing the ratio of ROI1 by ROI2. For the movement analysis in the *cals3-2d* mutant, the ROIs were selected by morphology, besides the outer most root cap cells, three layers of cells near the root tip were selected as ROI1, cells within 5 layers of epidermal cells were selected as the ROI2 region (Fig. S1). For imaging of embryos, ovules were isolated and mounted in a 4% paraformaldehyde/5% glycerol/PBS solution including 1.5% SCRI Renaissance Stain 2200 (R2200; Renaissance Chemicals) for counterstaining of embryos. After applying the coverslip, the embryos were squeezed out of the ovules, and R2200 and GFP fluorescence were visualized by excitation at 405 and 488 nm and detection between 430 to 470 and 500 to 535 nm, respectively. All confocal imaging was performed on a Leica SP5 II system equipped with hybrid detectors. For differential interference contrast (DIC) microscopy, dissected ovules were mounted in chloral hydrate solution (chloral hydrate, water, and glycerol 8:3:1, W/V/V). After incubating overnight at 4° C, samples were investigated with a Leica DMR microscope equipped with DIC optics.

### Phylogenetic and sequences comparison analysis

The phylogenetic relationship of selected bHLH family proteins were analysed by Clustal Omega (http://www.ebi.ac.uk/Tools/msa/clustalo/). The protein sequences were obtained from TAIR website (https://www.arabidopsis.org) with the following AGI numbers: AT1G19850 (ARF5/MONOPTEROS), AT2G47270 (bHLH151), AT3G25710 (bHLH32/TMO5), AT2G31215 (bHLH138), AT1G74500 (bHLH135/TMO7), AT2G41130 (bHLH106/TMO5L2), At1g64625 (bHLH157), AT3G06590 (bHLH148/AIF2), AT2G43060 (bHLH158), AT1G68810 (bHLH30/TMO5L1), AT1G09250 (bHLH149/AIF4), AT3G28857 (bHLH166/TMO7L4), AT3G05800 (bHLH150/AIF1), AT3G47710 (bHLH161/TMO7L1), AT3G17100 (bHLH147/AIF3), AT2G31280 (bHLH155), AT5G39860 (bHLH136/TMO7L3), and AT3G56770 (bHLH107/TMO5L3).

### Quantitative RT-PCR Analysis

Total RNA of Arabidopsis seedling roots was extracted with the RNeasy kit (QIAGEN). Poly(dT) cDNA was prepared from 1 μg of total RNA with an iScript cDNA Synthesis Kit (Biorad) and analyzed on a CFX384 Real-Time PCR detection system (BioRad) with iQ SYBR Green Supermix (BioRad) according to the manufacturer’s instructions. Primer pairs were designed with the Beacon Designer 7.0 (Premier Biosoft International). All individual reactions were done in triplicate with three biological replicates. Data were analysed with qBase (Hellemans et al., 2007). Expression levels were normalized to *ACTIN2* (AT3G18780). The oligo-nucleotide sequences are listed in Table S1.

### Statistical analysis

For relative intensity ratio comparison, we took confocal images from homozygous T3 lines (for p*TMO7*::3nGFP, p*TMO7*::TMO7-GFP, and p*TMO7*::TMO7-3GFP, *pTMO7::*TMO7-GFP/*clas3-2d, n*=19, 19, 18, and 13 respectively), homozygous T2 lines (for TMO7 likes: as least 6 independent lines and 18 to 25 images in total; for TMO7 linker-scanning mutants: at least 10 independent lines (beside *m10*, 5 independent lines, 11 images) and 15 images in total; for NLS and NES analysis: at 6 independent lines, 17 and 19 images; for TMO7-S39A and S42A, *n*= 6 and 7 independent lines, 13 and 16 images), and T1 lines (for bHLH138/151-GFP, *n*= 7 and 13; bHLH138-M5, -M8/9, -M5/8/9, *n*= 10, 7, and 4; bHLH151-M5, -M8/9, - M5/8/9, *n*=10, 15, and 4; for TMO7-S39,42A, *n*=7 independent lines, respectively). Images were taken and the ROI1/ROI2 ratio were calculated as described above. The data were box-plotted and analysed by one way ANOVA between every two data sets, the same label was give when p>0.05. Note that we use the same p*TMO7*::3nGFP, p*TMO7*::TMO7-GFP, and p*TMO7*::TMO7-3GFP data set as references in all figures and p*TMO7*::bHLH138- and bHLH151-GFP data set in Fig. 4 as references for clarity.

For CRISPR *tmo7* mutant root length analysis, seedling images were taken by scanning 7 days after germinating on 1/2MS-agar plate. The root length was analysed by ImageJ (https://imagej.net). Student t-test was used to analyse the significance.

### CRISPR-Cas9 tmo7 mutant generation

*tmo7* CRISPR constructs were designed as previously descripted (Tsutsui and Higashiyama, 2016) with minor modification. The short guidance RNA (sgRNA) sequence was designed in the reverse primer to amplify the U6 promoter and the forward primer to amplify the sgRNA scaffold. The two fragments were fused together using the sgRNA sequence as the overlapping region and amplified with primers with adaptor sequence complementary to pKIR1.1 vector. The fused fragment was further integrated with the pKIR1.1 by SLiCE cloning method (Zhang et al., 2014). After transformation, the red T1 Arabidopsis seeds were selected under the Leica M205 FA microscope equipped with epifluorescence. The T1 inflorescence apices were collected for genotyping (primers are listed in Table S1). The sequencing results were analysed using web-tool TIDE (https://tide.nki.nl). The T2 generation of the transformants with high genome modification probability were harvested and non-fluorescent seeds were grown for genotyping analysis. Homozygous T3 were selected for embryonic and rootless phenotype analyses.

## Acknowledgement

We thank Ykä Helariutta for generously sharing the *cals3-2d* mutant, Maritza van Dop, Colette ten Hove, Margo Smit, Prasad Vadepalli and Jos Wendrich for constructive feedback on the manuscript and members of the Weijers group for fruitful discussions.

## Author contributions

K.J.L., B.D.R and D.W. designed the experiments. H.v.M. and B.D.R. performed the *cals3-2d* rescue analysis. K.J.L. performed the rest of experiments. K.J.L and D.W wrote the manuscript with input from B.D.R˙.

